# Human Papillomavirus Infection Blunts STING Signaling and Alters Downstream Response

**DOI:** 10.1101/2025.11.06.686957

**Authors:** Isabelle G. Tobey, David P. Kosanke, Kelly M. King, Brittany L. Uhlorn, Advait Jeevanandam, Samuel K. Campos, Koenraad Van Doorslaer

## Abstract

The cGAS/STING pathway is an important innate immune pathway that senses and responds to foreign DNA or damaged host DNA. cGAS recognizes DNA and generates the second messenger cGAMP which activates STING. STING then creates a platform for IRF3 to be phosphorylated before pIRF3 induces a type I interferon response resulting in transcription of anti-viral genes. We examined how HPV18 infection modulates cGAS/STING activation following DNA stimulation in primary human foreskin keratinocytes from three different donors. Following exogenous activation, HPV(+) cells produced higher levels of cGAMP compared to patient-matched HPV(-) cells yet phosphorylation of STING and IRF3 was reduced in the HPV(+) cells. The reduced STING and IRF3 activation corresponded with a selective dampening of type I interferon driven antiviral genes and pro-inflammatory cytokines in HPV(+) cells. Simultaneously, HPV(+) cells had a baseline increase in genes associated with epithelial proliferation and skin development, which remained elevated following cGAMP treatment. We demonstrate that both E6 and E7 are required and sufficient to drive increased cGAMP levels and attenuate STING activation. Our results characterize cGAS/STING pathway activation across time in HPV positive and negative cells and demonstrate that the HPV oncogenes cooperate to suppress early STING activation despite higher levels of cGAMP in these HPV(+) cells. These findings indicate that HPV modulates cGAS/STING signaling to blunt antiviral defenses without impacting other arms of the cGAS/STING pathway response.

**Importance:** In this study, we used primary human keratinocytes from multiple donors to track cGAS/STING activity over time following stimulation. We found that although HPV(+) cells produced more cGAMP, these cells showed reduced early activation of STING and IRF3. Downstream of cGAS/STING, many antiviral genes were suppressed by HPV however, epithelial proliferation-related genes had an HPV-mediated increase in baseline expression which remained higher than uninfected cells following cGAMP stimulation. We identified the HPV oncogenes E6 and E7 work together to enhance cGAMP levels and suppress parts of the downstream STING signaling response. These findings reveal that HPV attenuates cGAS/STING signaling and downstream events, selectively suppressing antiviral genes without inhibiting other parts of the pathway. Our results provide new insights into how HPV modulates innate immune responses.

## Introduction

Human papillomaviruses (HPVs) are the most common sexually transmitted infection (Chesson et al., 2014), the majority of which are cleared by the immune system within two years (D’Souza et al., 2024). In order to maintain a persistent infection, HPV must evade detection and elimination by the immune system. Scagnolari et al. has shown that several components of the innate immune system are critical for eliminating persistent papillomavirus infection in the murine genital tract (Scagnolari et al., 2020). Infection of immunocompetent C57BL/6J mice with mouse papillomavirus MmuPV1 resulted in transient infections that were quickly cleared by the murine immune system. In contrast, MmuPV1 persisted for four months after initial infection in mice genetically deficient for the IL-1 receptor, the Toll Like Receptor adaptor MyD88, or STAT1. STING knockout resulted in varied but detectable MmuPV1 transcripts at four months, whereas virus was undetectable in wild type C57BL/6J mice, indicating that the cGAS/STING pathway may play an important role in counteracting persistent papillomavirus infection.

cGAS senses double-stranded DNA and generates the second messenger cGAMP. While cGAS was originally thought to function exclusively in the cytosol, it is now known to localize predominantly in the nucleus where it is tethered to chromatin in an inactive state, preventing binding of self-DNA and autoreactivity (Chen et al., 2020; Liu et al., 2018; Zhao et al., 2020). In the cytosol, following DNA-binding, cGAS generates the cyclic dinucleotide cGAMP, which activates STING (Ablasser et al., 2013; Diner et al., 2013; Gao et al., 2013), causing STING to translocate from the ER to the Golgi (Gui et al., 2019; Ishikawa et al., 2009; Mukai et al., 2016; Saitoh et al., 2009). In the Golgi, STING oligomerizes, recruits TANK binding kinase 1 (TBK1) and its homolog IKKε (Abe & Barber, 2014; Mukai et al., 2016), which ultimately results in the phosphorylation of IRF3 (Chen et al., 2016; Fitzgerald et al., 2003; Ishikawa & Barber, 2008; Sun et al., 2009; Tanaka & Chen, 2012; Zhong et al., 2008) and the NF-κB subunit p65 (Balka et al., 2020; Karin, 1999; Liu et al., 2017). phospho-NF-κB and phospho-IRF3 (pIRF3) enter the nucleus to initiate transcription of type I interferon (IFN) and pro-inflammatory cytokines (Balka et al., 2020; Chen et al., 2016; Hopfner & Hornung, 2020), which are capable of clearing HPV infections (Samuel, 2001; Shi et al., 2018; Terenzi et al., 2008).

HPV has been found to downregulate many parts of the cGAS/STING pathway (Albertini et al., 2018; Bortnik et al., 2021; Gusho & Laimins, 2022; Lau et al., 2015; Lo Cigno et al., 2020; Luo et al., 2020; Miyauchi et al., 2023; Saulters et al., 2024). While data suggests that HPV suppresses STING activity, there is a high degree of variability in the mechanisms of HPV suppression identified. Several researchers showed, by overexpressing STING and E7, that the LxCxE motif in E7 may be capable of directly binding STING (Lau et al., 2015; Shaikh et al., 2019). However, other researchers were unable to immunoprecipitate STING with endogenous HPV16 E7, indicating the two proteins may not be directly interacting under physiological conditions (Luo et al., 2020).

Similarly, there is conflicting data on whether HPV gene expression increases cGAS/STING protein levels or impacts protein phosphorylation/activation. Albertini et al. found that HPV positive cells had reduced steady state cGAS and STING protein levels both at baseline and following stimulation with the DNA analog poly(dA:dT) (Albertini et al., 2018). This decrease in cGAS and STING was corroborated by a study that showed knockout of E6 and E7 resulted in a significant increase in cGAS and STING mRNA (Lo Cigno et al., 2020). Conversely, Gusho and Laimins found that HPV16 or HPV31 positive keratinocytes had significantly more cGAS protein at baseline. They also saw that HPV positive cells had increased pathway activation (more phospho-STING and phospho-IRF3) after four hours of poly(dA:dT) or cGAMP stimulation compared to HPV negative cells (Gusho & Laimins, 2022). Still others showed that HPV positive cells had robust STING phosphorylation (i.e., activation) six hours post STING agonist stimulation but saw no downstream activation of TBK1 or IRF3, nor any HPV-mediated impact on cGAS, TBK1, or IRF3 protein levels (Saulters et al., 2024). Taken together, the above findings highlight several potential mechanisms by which HPV gene expression is modulating the cGAS/STING pathway.

Given that there is evidence that the cGAS/STING pathway may counteract the establishment of a persistent infection (Scagnolari et al., 2020), we sought to examine how HPV modulates pathway activation following acute stimulation of the cGAS/STING pathway in HPV positive and negative donor matched keratinocytes. We found that despite causing an increase in cGAMP, HPV18 positive (HPV18(+)) cells had delayed and reduced activation of STING and IRF3. The modest suppression of pathway activation corresponded with a selective decrease of antiviral type I interferon-driven (IFN-driven) genes and early cytokine signaling. In addition to the decrease in antiviral genes, there was an increase in epithelial proliferation and skin development genes that HPV upregulated at baseline and remained elevated following cGAMP treatment. Finally, we found that E6 and E7 play a synergistic role in muting STING and increasing cGAMP levels as co-expression of both oncogenes was required for the observed phenotypes.

## Materials & Methods

### Tissue Culture

Murine J2-3T3 fibroblasts were grown in high glucose DMEM supplemented with 10% NCS, 1% penicillin/streptomycin and 1% L-glutamine. Cells were maintained at 37°C with 5% CO2. Primary human foreskin keratinocytes (HFKs) were derived from foreskin samples as previously described (Van Doorslaer et al., 2016). HFKs labeled donors 1, 2, and 3 correspond to HFK12, 13, 17, respectively. Keratinocytes were passaged in Rheinwald-Green medium (3:1 Ham’s F12 (GIBCO 11765-054) / 4.5 µg/ml glucose Dulbecco’s modified Eagle’s medium (GIBCO 11960-044), 5% fetal bovine serum (Millipore Sigma F8067-500ml lot #16A328), 0.4 µg/ml hydrocortisone (Millipore Sigma H-4001), 8.4 ng/ml cholera toxin (Millipore Sigma 227036), 24 µg/ml adenine (Millipore Sigma A-2786), 6 µg/ml insulin (Millipore Sigma I1882), 10ng/ml epidermal growth factor (Thermo PHG0311), 4 mM L-glutamine (Thermo 25030-149), 50 µg/ml Penicillin/Streptomycin (Thermo 15140-148)). The Rheinwald-Green media was further supplemented with 10 μM Y-27632 dihydrochloride (Chemdea CD0141) (Chapman et al., 2014) when culturing HPV negative primary keratinocytes. Cells were maintained at 37°C with 10% CO_2_. Y-27632 dihydrochloride was omitted when HFKs were plated for use in experiments.

### HPV18 Cell Line Generation

HPV18 genomes were introduced into HFKs by electroporation, as previously described (Coursey & McBride, 2021). Briefly, circular HPV18 genomes were purified and electroporated into early-passage HFKs, which were then plated in Rheinwald-Green medium and cultured as described above, Y-27632 dihydrochloride. HPV18 positive status was confirmed by Southern blot.

### Nucleic Acid Transfections

HFKs were plated, without J2 fibroblast feeders, at 100,000 cells per well in 24-well plate or 600,000 cells per well in a 6-well plate. Cells were transfected with 500 ng or 2.5μg endotoxin-free pGL3 per well for 24-well or 6-well plates, respectively, using Lipofectamine 2000 (ThermoFisher 11668) in OptiMEM (Life Technologies). At various timepoints post-transfection, keratinocytes were washed with PBS and lysates collected.

### Generation of HPV18 E6/E7 Overexpression Constructs

To evaluate the individual and combined effects of HPV18 oncogenes on innate immune signaling, expression constructs for E6, E7, E6/E7, and an empty vector control were generated. The HPV18 E6 and E7 open reading frames were cloned into the retroviral vector pLSXN, as previously described (Halbert et al., 1991). To selectively express E6 or E7, translation termination linkers (TTL) were inserted early in either the E6 or E7 open reading frame, based on a previous design for HPV16 (Brimer & Vande Pol, 2022). The resulting sequences were synthesized by Twist Bioscience and cloned into the retroviral vector pLSXN.

### Retrovirus Production and Keratinocyte Transduction

HEK293T cells were maintained in DMEM supplemented with 10% FBS, 2 mM L-glutamine, and 100 U/mL penicillin-streptomycin. For retrovirus production, 293T cells were seeded at 1 × 10⁶ cells per 10 cm dish 48 hours prior to transfection. Cells were transfected with the following plasmids 5 μg pVSV-G, 2 μg CMV-tat, 10 μg MLV gag/pol, and 15 μg pLxSN retroviral constructs as described (Bartz & Vodicka, 1997). DNA was transfected using polyethylenimine (PEI)–DNA complexes prepared in Opti-MEM. Six hours post-transfection, transfection media was supplemented with complete HEK293T media, which was replaced 12 hours later with keratinocyte-specific recovery medium (Rheinwald-Green medium (3:1 Ham’s F12 (GIBCO 11765-054) / 4.5 µg/ml glucose Dulbecco’s modified Eagle’s medium (GIBCO 11960-044), 10% fetal bovine serum (Millipore Sigma F8067-500ml lot #16A328), 0.4 µg/ml hydrocortisone (Millipore Sigma H-4001), 8.4 ng/ml cholera toxin (Millipore Sigma 227036), 24 µg/ml adenine (Millipore Sigma A-2786), 6 µg/ml insulin (Millipore Sigma I1882), 4 mM L-glutamine (Thermo 25030-149)). Retroviral supernatant was harvested 48 hours after media change, filtered through a 0.45 µm filter, and supplemented with 10 µg/mL polybrene. J2 feeders were removed with versene as previously described (Coursey & McBride, 2021), and retrovirus was added to cells. After 16 hours, viral media was replaced with HFK passaging media containing irradiated J2 feeder cells and Y27632. 48 hours post-transduction, cells were selected with 400 μg/mL geneticin for 48 hours followed by maintenance selection at 200μg/mL G418. Retroviral expression was confirmed by western blotting using antibodies for p16^INK4A^ (Santa Cruz 56330 clone JC8) and PTPN14 (Cell Signaling 13808 clone D5T6Y), key markers of E6 and E7 expression.

### SDS-PAGE and Western Blotting

For western blot analysis, cells were lysed in 1x high SDS RIPA protein lysis buffer (50 mM Tris-HCl pH 8.0, 150 mM NaCl, 1% NP40, 0.5% sodium deoxycholate, 0.5% SDS), supplemented with 1x protease inhibitor cocktail (Sigma P1860), 1mM PMSF and 1x PhosSTOP phosphatase inhibitor cocktail (Roche 04906845001). Genomic DNA was sheared using a QIAshredder (Qiagen 79656) and samples were stored at −80°C until gel electrophoresis.

Total protein lysates were quantified using BCA assays (Thermofisher 23225). 20μg protein was separated on 10% or 12.5% polyacrylamide gels and transferred onto a 0.45 μm nitrocellulose membrane overnight. Blots were blocked in 5% non-fat powdered milk dissolved in Tris-buffered saline containing 0.1% Tween (TBST) for 1 hour at room temperature. Blots were probed with the following primary antibodies overnight at 4°C in 1% milk/TBST: rabbit monoclonal anti-cGAS (Cell Signaling 15102, 1:250), mouse monoclonal anti-p53 (Santa Cruz 47698, 1:200), mouse monoclonal anti-IRF3 (Abcam 50772 1:200), rabbit monoclonal anti-phospho-IRF3 (S386, Cell Signaling 37829, 1:500), rabbit monoclonal anti-STING (Cell Signaling 13647, 1:500), rabbit monoclonal anti-phospho-STING (S366, Cell Signaling 19781, 1:250), rabbit monoclonal anti-αTubulin (Abcam EP1332Y 1:40,000), mouse monoclonal anti-(Tubulin (Cell Signaling 3873, 1:500), mouse monoclonal anti-p16^INK4A^ (Santa Cruz 56330, 1:200). Goat anti-rabbit DyLight 680 (Pierce 35568), goat anti-mouse DyLight 680 (Pierce 35518), goat anti-rabbit DyLight 800 (Pierce 535571) and goat anti-mouse DyLight 800 (Pierce 35521) were used as secondary antibodies at 1:10,000 in 1% milk/TBST for 1 hour at room temperature.

Blots were imaged using the Licor Odyssey Infrared Imaging System. Band intensities were quantified by densitometry using imageJ v2.14.0/1.54f (Schneider et al., 2012). Briefly, boxes were drawn around individual bands and local background intensity was subtracted from each band. Total STING, pSTING, total IRF3, and pIRF3 values were normalized to αTubulin and reported as arbitrary units. For multi-blot experiments (Figure 5) the same control lysate was run on each gel and densitometry was first normalized to αTubulin and then normalized to a control lane.

### 2’-3’ cGAMP ELISA

For cGAMP ELISA, cells were transfected as above, rinsed in PBS, and lysed in 1x low SDS RIPA lysis buffer (50 mM Tris-HCl pH 8.0, 150 mM NaCl, 1% NP40, 0.5% sodium deoxycholate, 0.1% SDS). Lysates were frozen at −80°C, thawed to 37°C to scrape the plates, and stored at −80°C until ELISA.

Lysates were pelleted at 10,000xg for 10 minutes at 4°C and the supernatant was used to quantify 2’-3’ cGAMP via ELISA (Cayman Chemical 501700) following the manufacturer’s protocol. 2’-3’ cGAMP concentrations were normalized by total protein concentration as determined by BCA assay (Thermofisher 23225) and results were plotted as pg cGAMP/μg total protein.

### Bulk RNA Sequencing and Analysis

Primary human foreskin keratinocytes (HFKs) with or without HPV18 genomes were treated with 25 µg/mL 2′3′-cGAMP (InvivoGen tlrl-nacga23) to directly activate STING. In addition to cGAMP, cells were also treated with either 1μg/mL IFNAR1-blocking antibody (PBL Assay Science 21370, clone MMHAR-3) or 1μg/mL IgG1 isotype control antibody (Invitrogen 14471485, clone P3.6.2.8.1) to distinguish IFN-dependent from IFN-independent signaling. RNA was harvested at 0, 3, 6, or 9 hours post-treatment. Untreated cells collected at 0 hours served as baseline controls. Cells were lysed using TRIzol Reagent (Invitrogen 15596026) and lysates were shipped to Novogene Co. (Sacramento, CA) for RNA extraction, library preparation, and sequencing on the Illumina NovaSeq X Plus platform, yielding >20 million paired-end reads (2 × 150 bp) per sample. Raw sequencing reads are deposited in the Sequence Read Archive (SRA) under accession numbers PRJNA1357338.

High-throughput sequencing data were initially processed using MultiQC (Ewels et al., 2016) for quality screening and the STAR aligner (v2.7.3a) (Dobin et al., 2013) for mapping reads to the human reference genome version GRCh38 and generation of gene-level read counts. Differential gene expression analysis of read counts, including library-size correction and statistical testing, was performed using DESeq2 (Love et al., 2014) in R (v4.4.2). Independent filtering was turned off to capture low expressing genes.

A likelihood ratio test (LRT) was used to identify genes that exhibited significant changes over time in response to cGAMP stimulation in HPV-negative, isotype-control treated cells. Genes with an adjusted p-values <0.05 were considered significantly responsive. To identify differentially expressed genes (DEGs) at each time, we performed pairwise Wald tests comparing each time point (3, 6, 9 hours) to baseline (0 hr). DEGs were defined as padj<0.05 and |log2FoldChange|≥ 0.5. In total, 1856 genes were significantly altered by cGAMP, including 831 upregulated and 1025 downregulated transcripts. To assess how HPV alters cGAMP-responsive gene expression, we analyzed DEGs between HPV-positive and HPV-negative samples for each time point (3, 6, 9hr) using the same thresholding as before. 2901 genes were upregulated in HPV(+) cells compared to HPV(-) and 2031 genes were suppressed.

We examined genes that were induced by cGAMP in the HPV(-) cells yet significantly up or downregulated in the HPV(+) cells by looking for overlaps between the datasets and identified 284 genes that were upregulated by HPV (Table S1) and 263 genes that were suppressed (Table S2). Additionally, there were 285 genes that were induced by cGAMP in HPV(-) cells but not significantly altered by HPV (Table S3). Unsupervised hierarchical clustering was used to determine trends in gene expression patterns, which identified 12 primary clusters, six upregulated by HPV and six suppressed (Table S4). To assess the contribution of type I IFN signaling to these clusters, we examined the gene expression patterns in the presence of an IFNAR-blocking antibody. Z-scored expression levels for HPV(+) and HPV(-) cells with and without IFNAR-blocking antibody were plotted for each cluster. Gene Ontology enrichment analysis for biological processes (GO:BP) was performed using the *clusterProfiler* package (Xu et al., 2024). Enriched terms were identified with *enrichGO* applying significance thresholds of p-value ≤ 0.1 and q-value ≤ 0.1. Networks of the top 5 enriched GO terms per cluster were visualized using the Kamada-Kawai layout with pie charts where the size indicates the number of genes in each node and the colors relate to the different clusters.

## Results

### HPV18 increases cGAMP levels following exogenous activation

To examine the interplay between HPV and the cGAS/STING pathway, we generated three sets of donor-matched HPV18 positive and negative cell lines from primary foreskin keratinocytes (Meyers et al., 1992; Sterling et al., 1990). Following establishment of the viral genome, we transfected these cells with double-stranded plasmid DNA to activate the cGAS/STING pathway (Uhlorn et al., 2020), and collected lysates every two hours for eight hours. Intracellular cGAMP levels were measured by ELISA. Without exogenous stimulation, cGAMP levels not consistently above the limit of detection (LOD). Following DNA transfection, both HPV18(+) and parental HPV(-) cells showed an increase in intracellular cGAMP levels over time (Fig. 1A). At six- and eight-hours post-transfection, HPV18(+) cells produced significantly more cGAMP than HPV negative cells. These data indicate that despite donor variability (Fig. 1B), HPV18 increases intracellular cGAMP upon DNA transfection, compared to uninfected cells.

**Figure 1:**
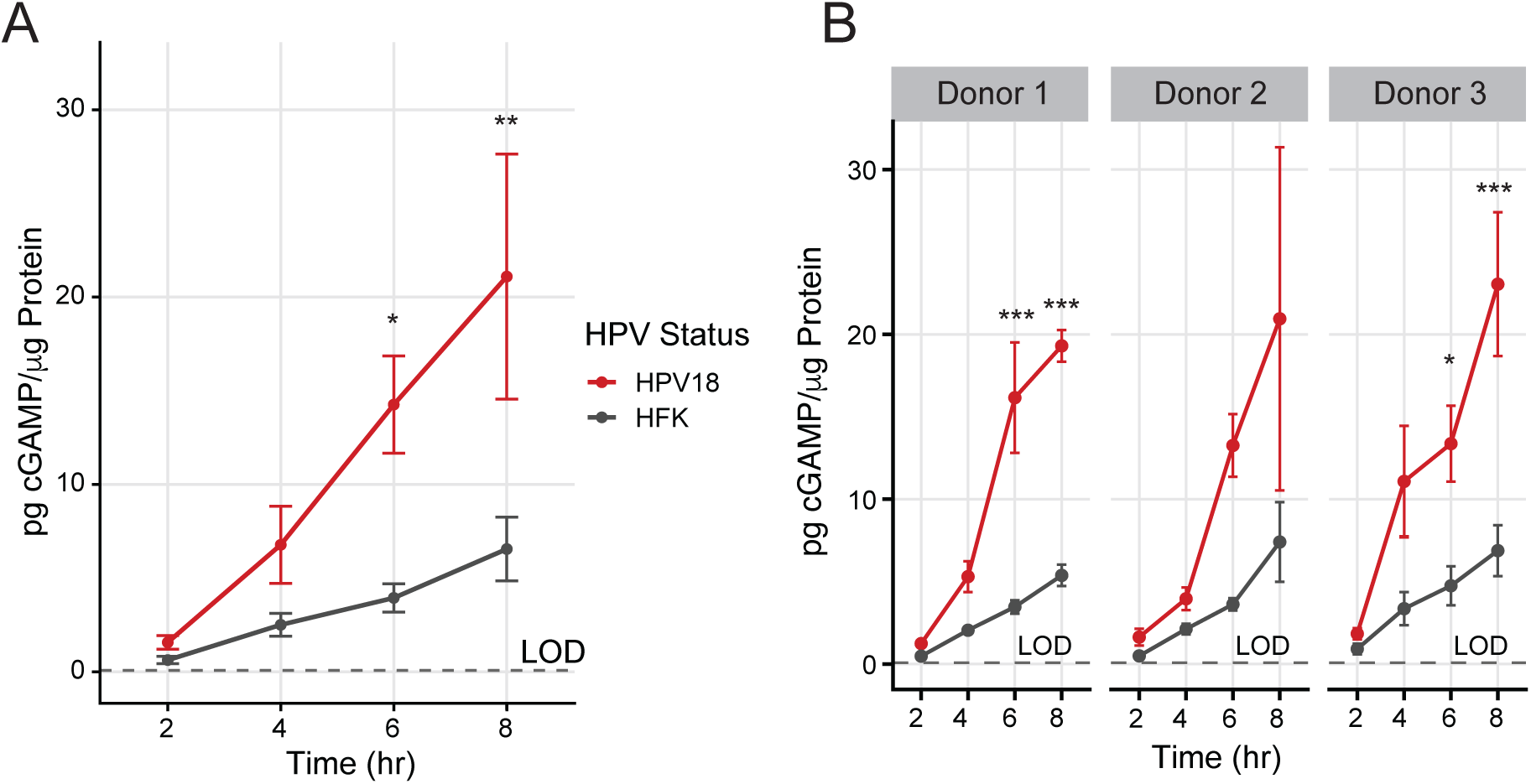
HPV18(+) cells produce more cGAMP than parental control cells. Quantification of cGAMP by ELISA following DNA stimulation of HPV(-) cells (black) and HPV18(+) cells (red). N=3 donors, 3 reps in each donor, average of the 9 replicates, (A) replicates separated by donor (B). Statistics for (A) were calculated using a linear mixed-effects model with “HPV” and “Time” as fixed effects and donor as a random effect to account for inter-donor variability. Statistics for (B) were calculated with a two-way ANOVA followed by a Dunnett’s post hoc test. * p<0.05; ** p<0.01; *** p<0.001Error bars for represent SEM, calculated by first averaging technical replicates (n=3 per donor) with propagated error, and then computing SEM across biological replicates (n=3 donors).

### HPV18 reduces the phosphorylation of STING and IRF3 downstream of cGAS activation

To examine whether the presence of HPV18 genome and/or the observed increase in cGAMP levels (Fig. 1) impact downstream cGAS/STING signaling, we again transfected cells with DNA. We detected and quantified STING, IRF3, and their activated (phosphorylated) forms by western blot (Fig. 2A). p16^INK4A^, induced by HPV, (Carozzi et al., 2008; McLaughlin-Drubin et al., 2013) was detected only in the HPV(+) cells. In the parental HPV(-) cells (Fig. 2B, black line), we observed an increase in the phosphorylation of STING following exogenous DNA stimulation. This shows that the transfected DNA is sensed by cGAS, leading to the production of cGAMP (Fig. 1) and the activation of STING. STING was rapidly phosphorylated following stimulation, peaking at two hours post transfection. By eight hours post stimulation, pSTING returned to baseline levels (p=0.17, 0hr vs 8hr). HPV18(+) cells followed similar kinetics, peaking at two hours and returning to baseline line by eight hours post stimulation (red line). HPV18(+) cells had a reduced pSTING response (Fig. 2B). While all three donor lines showed peak pSTING at two hours post stimulation, each donor had variability in the extent of STING activation and the degree of HPV-mediated suppression (Fig. 2C). All three donors showed strong initial induction between 0-1hrs, where the rate of increase in pSTING was significantly suppressed in HPV(+) cells (average slope = 11.6 ± 1.7) compared to HPV(-) cells (average slope = 23.0 ± 4.5, linear mixed-effect model (LMM) interaction p = 0.005), indicating STING activation was both slower and reduced in the HPV(+) cells. By four hours, we observed a reduction in pathway activation in all cell lines as pSTING levels began to decrease. This decrease was equivalent in HPV(+) and HPV(-) cell lines.

**Figure 2.**
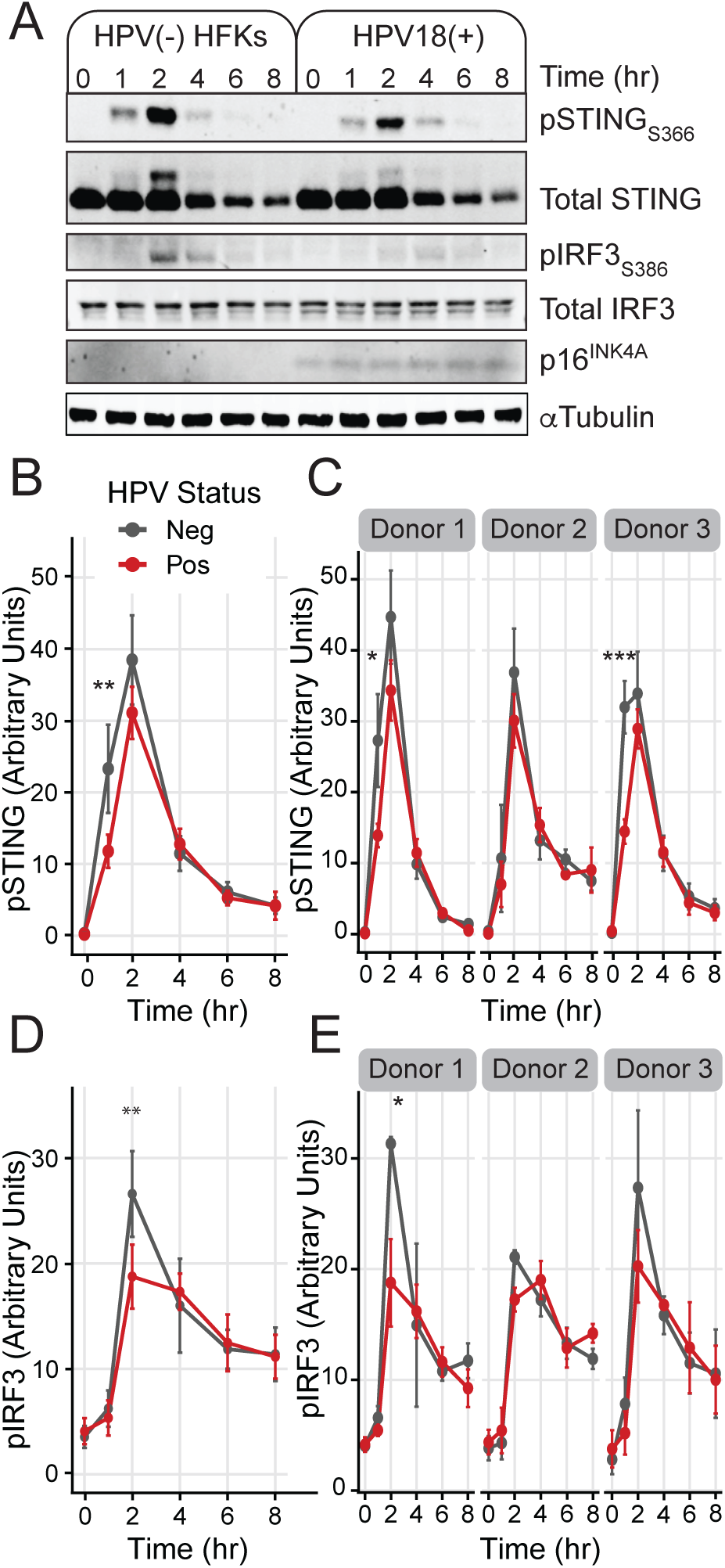
HPV decreases STING and IRF3 activation. (A) Western blot showing DNA-treated HPV18(+) and parental cells. Quantification of (B) pSTING, average of the 9 replicates (C) pSTING, separated by donor (D) pIRF3 western blots, average of the 9 replicates, (E) pIRF3, separated by donor. N=3 donors, 3 replicates in each. Statistics for the average of 9 (B and D) were calculated using a linear mixed-effects model with “HPV” and “Time” as fixed effects and donor as a random effect to account for inter-donor variability. Statistics for each cell line (C and E) were calculated with a two-way ANOVA followed by a Dunnett’s post hoc test. * p < 0.05; ** p<0.01; *** p<0.001. Error bars represent SEM, calculated by first averaging technical replicates (n=3 per donor) with propagated error, and then computing SEM across biological replicates (n=3 donors). Slope was calculated by fitting linear regressions within each time interval, and the values were subsequently analyzed using a linear mixed-effects model.

Further downstream the canonical cGAS-STING pathway, IRF3 phosphorylation was also muted in HPV18(+) cells following DNA stimulation (Fig. 2A). In HPV(-) cells, we observed a significant increase in pIRF3 compared to unstimulated cells (Fig. 2C) which peaked two hours post stimulation (p<0.001, 0hr vs 2hrs). By eight hours post transfection, the amount of pIRF3 remained significantly above baseline (p<0.001) and did not reset to pre-transfection levels. In HPV18(+) cells, pIRF3 levels peaked at two hours, but were significantly lower than in uninfected cells (p<0.01) and remained suppressed through four hours. As before, there was variability in the extent of response and HPV suppression within the three donors (Fig. 2D). All three donors had minimal activation of IRF3 at one hour post stimulation but strong induction between 1-2hrs. In the HPV(+) cells, the rate of increase in pIRF3 was slower from 1-2hrs (average slope = 13.4 ± 1.8) than in the HPV(-) cells (average slope = 20.4 ± 2.7, LMM interaction p=0.02), suggesting that HPV delays and suppresses pathway activation.

These data indicate that despite donor-to-donor variability, HPV18 delays and reduces, but does not fully block, STING and IRF3 activation following exogenous DNA transfection. Notably, the delayed pSTING and pIRF3 response occurred despite the increase in cGAMP in HPV18(+) cells (Fig. 1), which is expected to activate STING.

### HPV reshapes gene expression downstream of STING and IRF3

To assess the transcriptional consequences of reduced STING/IRF3 activation, we stimulated donor lines and patient-matched HPV18(+) cells with 25μg/mL cGAMP and measured RNA expression over time. In HPV(-) cells, cGAMP significantly induced a combined 831 genes at three, six, or nine hours post-cGAMP-treatment compared to untreated cells (0 hours; p-adjusted <0.05, and fold change >0.5). Among these, 263 genes that were normally upregulated by cGAMP were suppressed in HPV(+) cells (Table S1), while 284 were significantly increased by HPV compared to uninfected cells (Table S2), and 285 were not significantly changed by the presence of HPV (Fig. 3A, Table S3). Most of the 284 increased genes were elevated at baseline in the HPV(+) cells. Since IRF3 activation leads to a type I IFN response (Chen et al., 2016; Hopfner & Hornung, 2020), we also treated cells with an Interferon-Alpha/Beta Receptor (IFNAR) blocking antibody or an IgG isotype control.

**Figure 3.**
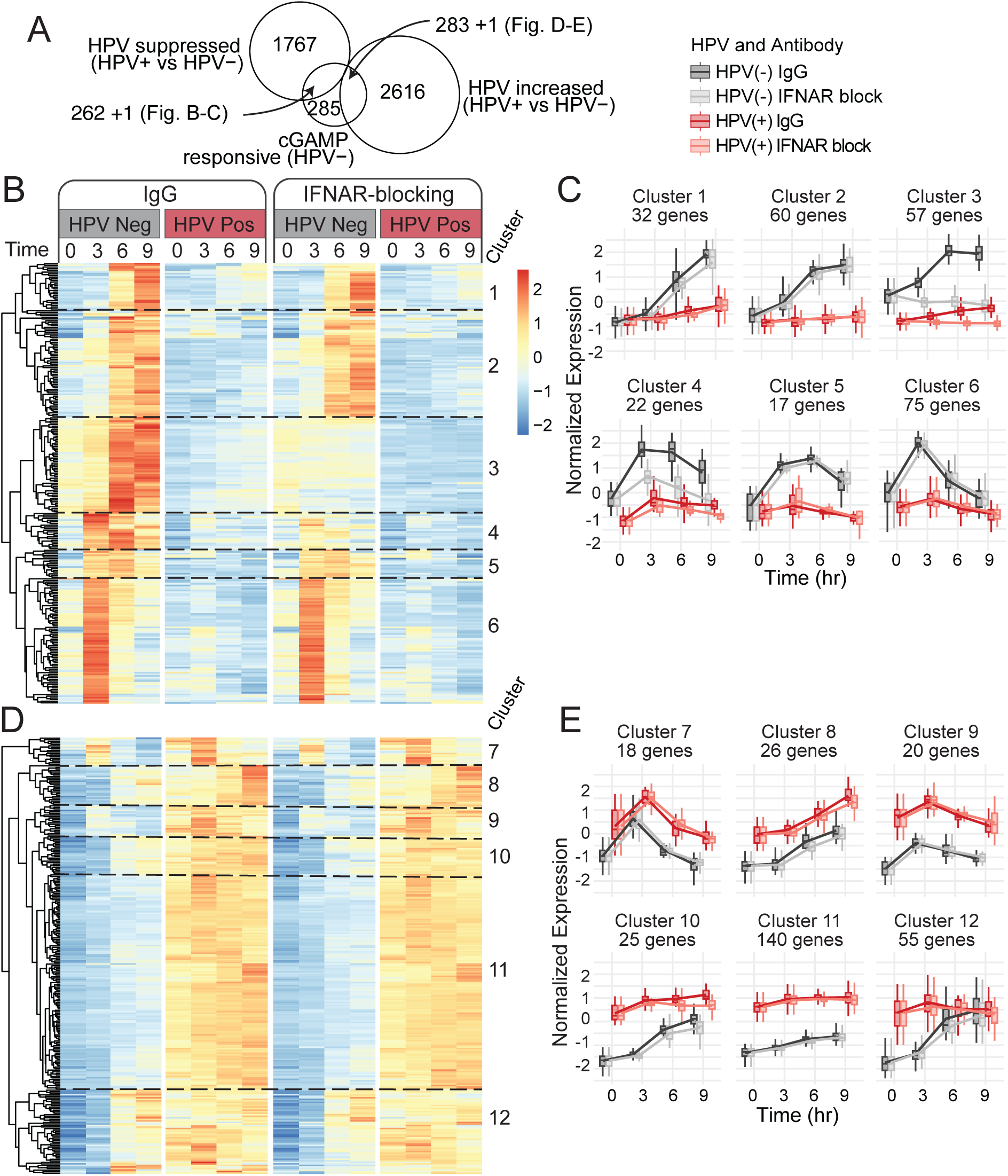
Identification of cGAMP-responsive genes modulated by HPV. (A) Euler plot showing the overlap between 831 cGAMP-responsive genes identified in HPV(-) cells, 2031 HPV suppressed genes, and 2900 HPV increased genes. DEGs defined as padj < 0.05, |log2FC| ≥ 0.5. (B) Z-scored heatmap of the 263 DEGs shared between the cGAMP responsive genes and the HPV suppressed genes. (D) Z-scored heatmap of the 284 DEGs shared between the cGAMP responsive genes and the HPV increased genes. Heatmaps are divided into clusters based on independent hierarchical clustering. (C, E) Z-scored expression profile following IgG or IFNAR-blocking antibody treatment for genes in panel B and E, respectively.

Hierarchical clustering organized these genes into 12 clusters with distinct dynamics (Fig. 3B, D, Table S4). One gene (GBP6) was included in both lists as it was significantly suppressed in HPV(+) cells at three hours post stimulation and significantly increased at nine hours. Clusters 1–6 represented the 263 genes induced by cGAMP in HPV(-) cells but attenuated in HPV(+) (Fig. 3B-C). Clusters 3 and 4 contained classic interferon-stimulated and antiviral genes, including CXCL10/11, OAS1, and IFIT1/3. HPV strongly reduced gene expression within these clusters. Clusters 3 and 4 were also sensitive to IFNAR blockade, suggesting that their activation partially depends on secondary type I IFN signaling. Cluster 6 showed transient induction in HPV(-) cells and was largely unaffected by IFNAR inhibition, suggesting more direct regulation by cGAS/STING. Activation of the genes in cluster 6 was fully blunted in HPV18(+) cells. Cluster 6 contained well known inflammatory cytokines including IL1A, IL1B, IL20, and IFNL1. Clusters 1, 2, and 5 included cell-cycle–associated genes, which were consistently blunted in HPV(+) cells and largely insensitive to IFNAR blockade. Clusters 1–6 indicate that HPV suppresses both IFN-dependent antiviral transcriptional programs and IFN-independent regulation of the cell cycle following activation of the cGAS/STING pathway.

Clusters 7–12 are defined by the 284 genes increased by cGAMP in uninfected cells that show significant elevation in the HPV(+) cells at one or more time points relative to HPV(-) cells (Fig. 3D-E). These clusters are upregulated by HPV at baseline (i.e., prior to cGAMP addition) when compared to parental cells and gene expression remained higher after cGAMP stimulation. Clusters 7, 8, 9, and 11 included genes that were largely independent of IFNAR signaling. In contrast, clusters 10 and 12 showed modest sensitivity to IFNAR blockade, suggesting a partial contribution of type I IFN signaling to these gene expression dynamics. With the exception of clusters 10 and 12, all observed genes follow the same dynamics in HPV18(+) and control cells, suggesting that HPV is not altering the transcriptional activation of these genes in response to cGAMP addition. Genes in clusters 10 and 12 increase following cGAMP treatment of parental cells, but not in HPV18(+) cells which indicates that these genes are no longer cGAMP responsive in HPV(+) cells.

To identify enriched biological processes across gene clusters, we performed gene ontology biological processes (GO:BP) enrichment analysis (Fig. 4) and plotted the top 5 GO:BP terms per cluster. Network mapping revealed that genes suppressed by HPV (clusters 1-6) were primarily enriched for cell cycle progression, antiviral response, and broad immune functions such as “response to molecule of bacterial origin”. In contrast, clusters 7-12 yielded fewer enriched terms, but uniquely highlighted pathways related to proliferation, skin development, salt response, and steroid biosynthesis. Despite being the largest cluster, cluster 11 had no significantly enriched GO:BP terms associated with it. Several GO:BP terms were enriched in gene clusters both upregulated and downregulated by HPV, such as “defense response to virus” which was primarily enriched in clusters 1-6 but also cluster 7. This suggests that even within antiviral biological processes, HPV may be selectively enhancing certain genes and “rewiring” the cGAMP response. The twelve clusters suggest that HPV does not abolish cGAS/STING signaling but instead reshapes it. HPV selectively blocks classical IFNAR-dependent genes and pro-inflammatory cytokines, while simultaneously amplifying proliferation and skin development programs that are largely unresponsive to cGAMP. This pattern is consistent with our data showing that HPV modestly blunts STING and IRF3 signaling without fully suppressing activation (Fig. 2).

**Figure 4.**
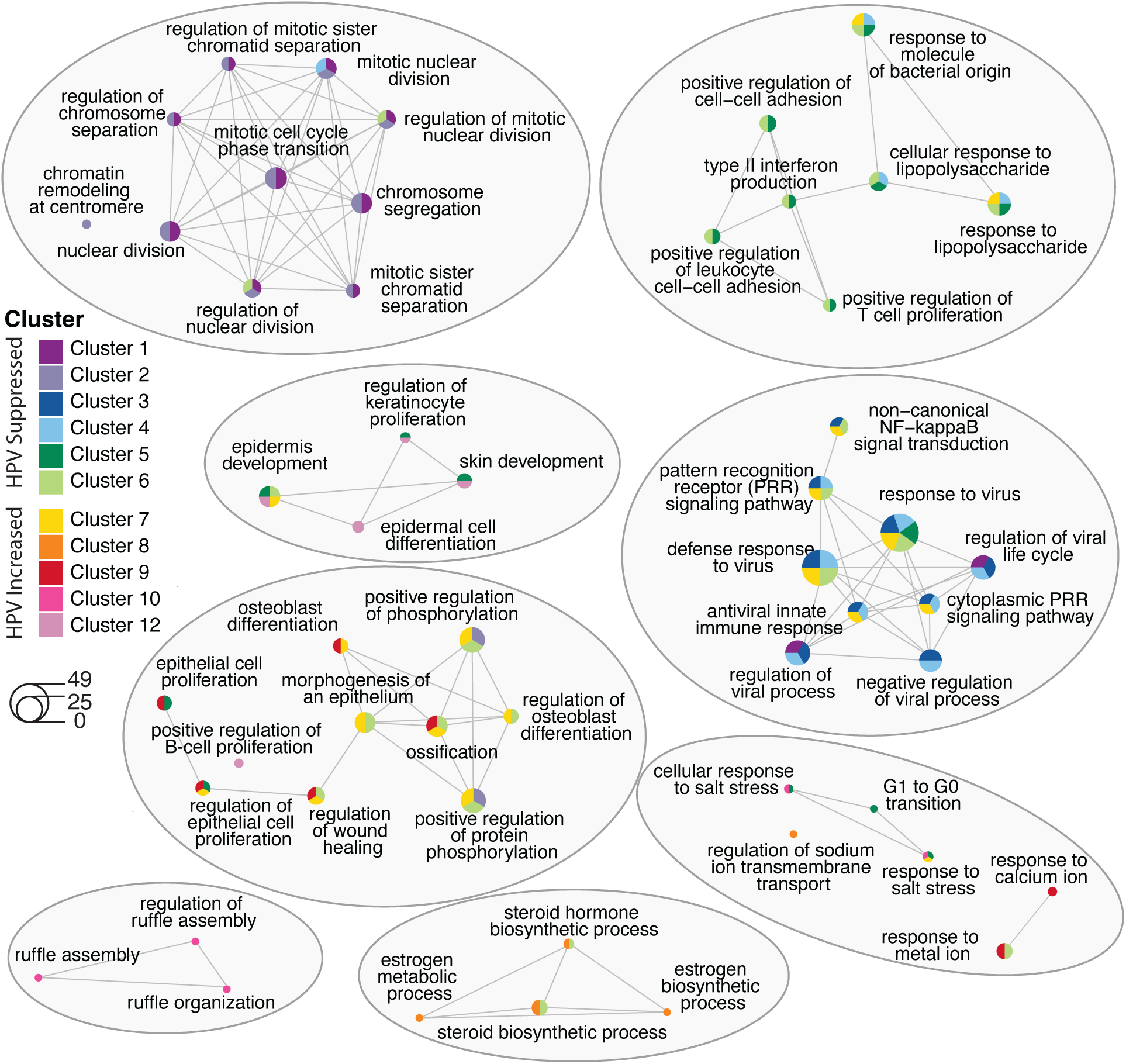
GO:BP enrichment network of HPV-regulated gene clusters. Network map of top 5 enriched biological processes identified per cluster. GO:BP analysis used a threshold of p ≤0.1 and q≤0.1. Each node represents an enriched GO:BP term with edges representing functional similarity between terms. Node size corresponds to the number of genes contributing to the term and colored pie charts indicate the relative contribution of each cluster. Network analysis was visualized using the force-directed Kamada-Kawai layout.

### HPV18 E6 and E7 are required and sufficient to increase cellular cGAMP concentration

To further understand how HPV differentially modulates the cellular cGAMP/STING response, we tested the contributions of the viral oncogenes, E6 and E7. We generated cell lines that contain viral genomes unable to express E6, E7, or both. These mutant genomes were generated in donor 1 (Fig. 1B, 2C, 2E) and in the background of the recombinant HPV18-Neo genome (Van Doorslaer et al., 2016) by inserting short translation termination linkers (TTL) into the respective open reading frames. These TTL insertions introduced a premature stop codon resulting in the functional deletion of the oncogenes without altering the expression of other viral genes. The successful deletion of specific viral proteins was confirmed via western blot (Fig. 5A). p53 and PTPN14 were used as markers for functional deletion of E6 and E7, respectively. We observed degradation of p53 when cells express E6 (compare lanes 2 and 4 to lanes 1, 3, and 5). As previously reported, expression of E7 stabilized p53 in cells not expressing E6, resulting in elevated p53 expression (lane 3; (Seavey et al., 1999)). Cells expressing E7 protein showed a degradation of PTPN14 (compare lanes 2 and 3 to lanes 1, 4, and 5).

**Figure 5.**
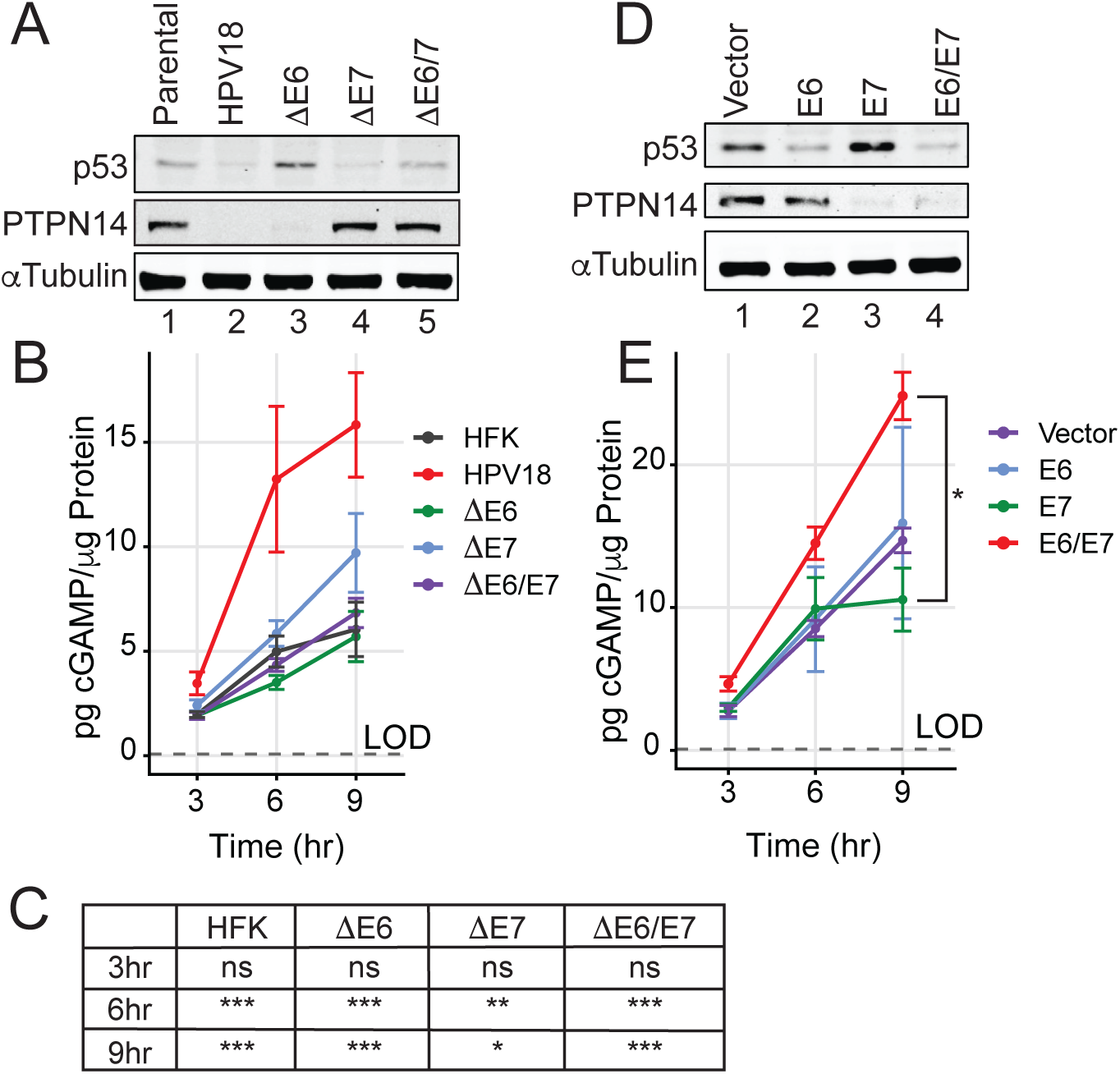
Both oncogenes are needed to impact cGAMP levels. (A) western blot analysis for functional markers p53 and PTPN14, N= 3 using TTL system (B) cGAMP ELISA following DNA stimulation, N=4. (C) p-values for cGAMP ELISA of each cell line relative to HPV18 at specified time points. (D) Western blot showing function expression of E6 and/or E7 in overexpression cell lines. N=3 (E) cGAMP ELISA following DNA transfection, N=3 Statistics for (C and E) were calculated with two-way ANOVA, followed by Dunnett’s multiple comparison test. * p < 0.05; **p < 0.01; *** p < 0.001. Error bars indicate SEM.

We transfected these cells with exogenous DNA and collected lysates at three-, six-, and nine-hours post stimulation. Cells with wildtype HPV18 genome expressing both E6 and E7 showed a statistically significant increase in intracellular cGAMP levels compared to all other cell lines at six- and nine-hours post DNA stimulation (Fig. 5B, C), suggesting that both E6 and E7 expression is needed for the stimulation-dependent increase in cGAMP in HPV(+) cells.

To determine whether E6 and/or E7 is sufficient to regulate the cGAS/STING pathway, we generated primary HFKs expressing HPV18 E6, E7, E6 and E7, or a negative vector control. To generate these vectors, we cloned a fragment of the HPV18 genome (nucleotide 826 to 1628) which encodes E6 and E7, into the pLSXN retroviral vector (Halbert et al., 1991), as previously described for HPV16 (Brimer & Vande Pol, 2022), allowing expression of E6 and E7 genes from polycistronic mRNAs. To express E6 and E7 individually, we inserted a TTL in the E7 or E6 open reading frame, respectively, as well as two separate TTLs into both ORFs to generate a control vector unable to express either viral protein. We then transduced these retroviral constructs into donor 1 cells and confirmed functional expression of viral oncogenes via western blot for p53 and PTPN14 (Fig. 5D). Cells expressing E6 (lanes 2 and 4) showed reduced p53 levels while those expressing E7 (lanes 3 and 4) showed decreased PTPN14 expression.

We transfected these cell lines with DNA and quantified cGAMP levels in these cells at three-, six-, and nine-hours post transfection (Fig. 5E). Cells expressing both E6 and E7 (red) had the highest levels of cGAMP. The increase in cGAMP was dependent on expressing both E6 and E7 as cell lines expressing individual viral proteins (blue or green curves) showed no significant change relative to vector containing cells (purple). These data suggest that co-expression of both E6 and E7 is sufficient and necessary for driving the increase in cGAMP levels following stimulation.

### HPV18 E6 and E7 cooperate to limit STING phosphorylation

We used the cell lines described in Figure 5 to test the impact of viral gene expression on the phosphorylation of STING. First, we transfected the HPV18-Neo (and derived TTL) genomes with DNA and quantified pSTING using western blot (Fig. 6A-C). HFKs maintaining wt HPV18(+) genomes showed reduction of pSTING compared to uninfected cells and cells unable to express E6/E7 (statistically significant at two hours post activation). Cells unable to express either E6 (green) or E7 (blue) alone did not reduce the pSTING levels to the same extent as the wildtype genome containing cells and were less activated compared to the parental control cells, indicating that both viral proteins may be involved in regulating the phosphorylation of STING.

**Figure 6.**
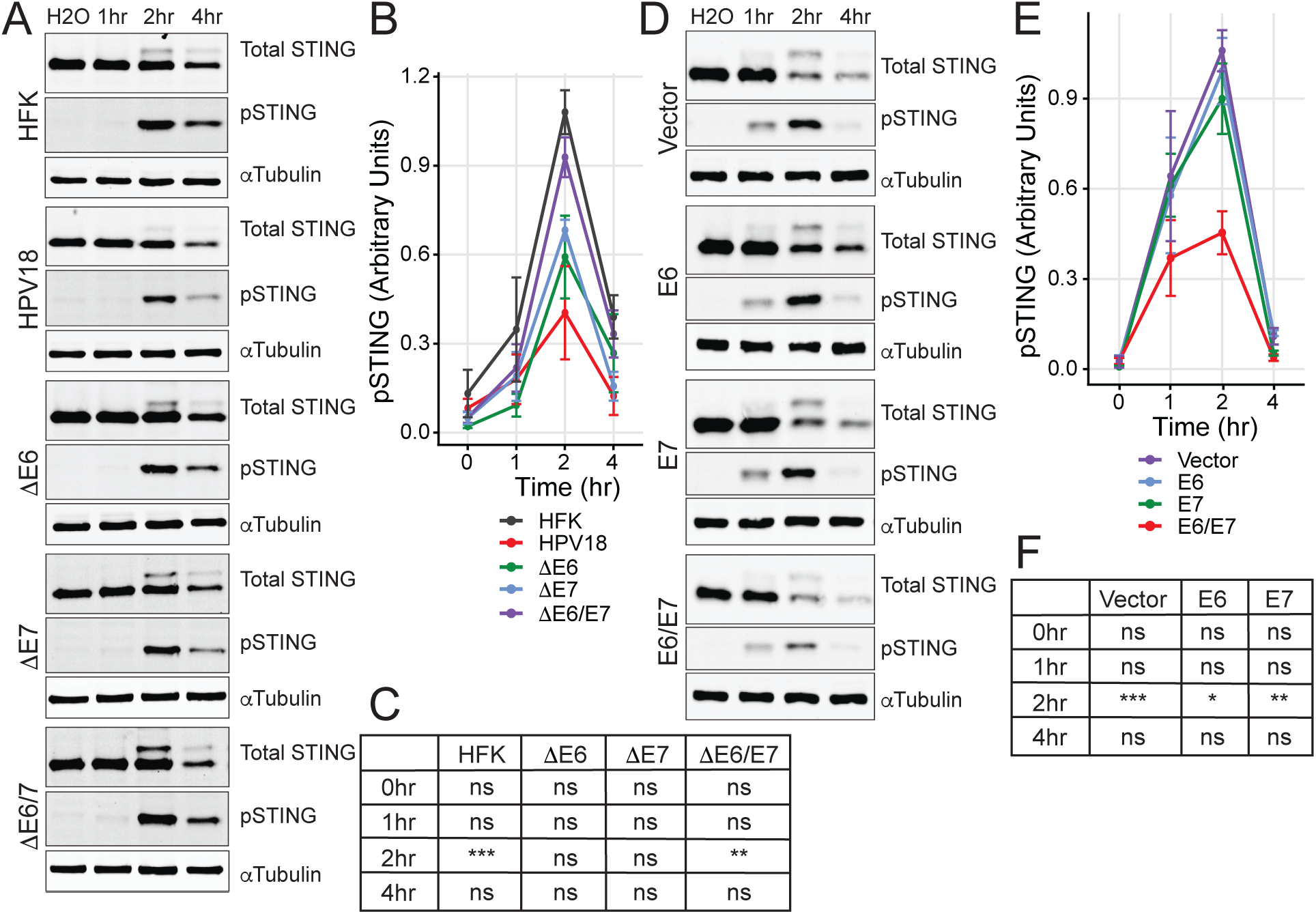
Both oncogenes are needed to impact STING activation. (A) Western blot for total STING, pSTING, and αTubulin for the TTL cell lines, N=3. (B) Densitometry of pSTING normalized to αTubulin and a loading control lysate run on all blots. (C) p-values for pSTING densitometry of each cell line relative to HPV18 cells at specified time points. (D) Western blot for total STING, pSTING, and αTubulin for the over expression cell lines, N=3. (E) Densitometry of pSTING normalized to αTubulin and a loading control lysate run on all blots. (F) p-values for pSTING densitometry of each cell line relative to E6/E7 expressing cells at specified time points. Statistics for (C and F) were calculated with two-way ANOVA, followed by Dunnett’s multiple comparison test. * p < 0.05; ** p < 0.01; *** p < 0.001. Error bars indicate SEM.

We then used the E6, E7, or E6/E7 expressing cell lines to further assess the role of the individual viral proteins. We quantified pSTING levels at one-, two-, and four-hours post transfection with exogenous DNA (Fig. 6D-F). As seen previously, pSTING levels peaked at two hours post-transfection. Cells expressing the control vector (purple) or individual proteins (E6 – blue or E7 – green) had equivalent pSTING levels in these experiments. Only cells that express both E6 and E7 (red) had a significantly reduced STING response upon exogenous DNA transfection.

Taken together, these data show that in cells with the full genome, expression of a single oncogene (either E6 or E7) is sufficient to partially reduce pSTING but both E6 and E7 are needed for full suppression. However, in the overexpression lines, both oncogenes are needed to achieve STING suppression.

## Discussion

The cGAS/STING pathway plays a central role sensing and responding to foreign or host DNA (Zheng et al., 2020). HPV has been found to induce genomic DNA damage (Duensing & Münger, 2002, 2004; Korzeniewski et al., 2011; Mehta & Laimins, 2018), increase R-loop formation (Templeton & Laimins, 2024), and release viral genomes into the cytosol (Bienkowska-Haba et al., 2023; Shin et al., 2025). Genomic damage, R-loops, and cytoplasmic viral DNA can be detected by the antiviral pattern recognition receptor cGAS, which results in the production of immune activating cytokines capable of clearing viral infections (Crossley et al., 2023; Durbin et al., 2000; Shi et al., 2018; Wieland et al., 2000; Zheng et al., 2020). Since there are several pathogen-associated molecular patterns HPV may be generating that can stimulate the cGAS/STING pathway, it is not surprising that there is mounting evidence indicating HPV suppresses cGAS/STING activation (Albertini et al., 2018; Bortnik et al., 2021; Gusho & Laimins, 2022; Lau et al., 2015; Lo Cigno et al., 2020; Miyauchi et al., 2023; Saulters et al., 2024; Shaikh et al., 2019). However, the prior research varies in regard to cell lines, HPV types, methods, and, ultimately, findings. For example, HPV has been reported to both decrease cGAS protein levels (Albertini et al., 2018; Lo Cigno et al., 2020; Tolbert et al., 2024) as well as increase levels (Gusho & Laimins, 2022; Luo et al., 2024). Similarly, while several studies report direct interactions between E7 and STING (Lau et al., 2015; Shaikh et al., 2019), others were unable to recapitulate this finding (Luo et al., 2020). Following stimulation, HPV positive cells have been shown to have decreased STING protein levels (Albertini et al., 2018), an E6- and E7-dependent decrease in STING mRNA (Lo Cigno et al., 2020) or have an increase in pSTING (Gusho & Laimins, 2022). In addition to E6 and E7 driven changes in cGAS/STING activation, recent studies have identified E5 may also be suppressing activation (Miyauchi et al., 2023). Together, this highlights the many ways by which HPV may be modulating cGAS/STING pathway activation.

In this study, we tracked how HPV affects cGAS/STING signaling, across time and in multiple donors. We found that HPV(+) HFKs have elevated cGAMP levels following DNA-stimulation compared to patient-matched HPV negative cells. Although baseline cGAMP levels were below the limit of detection, HPV(+) cells in all three donors consistently produced more intracellular cGAMP at all post-stimulation time points, with levels significantly higher at six- and eight-hours. We found that both E6 and E7 expression were required to drive increased cGAMP levels. In the TTL cell lines, the full HPV18 genome led to significantly higher levels of cGAMP at six- and nine-hours post stimulation compared to all other cell lines. Similarly, in the overexpression system, cells expressing both E6 and E7 consistently showed the highest cGAMP levels.

Despite containing cGAMP, HPV18(+) cells showed a significant reduction in early pSTING and pIRF3 responses to exogenous DNA stimulation, suggesting that HPV delays and suppresses the cGAS/STING response. While donor-specific differences were observed in both pathway activation and the extent of HPV-mediated suppression, all cell lines consistently showed peak IRF3 activation at two hours post stimulation, which corresponded to a subtle but consistent suppression of pIRF3 in HPV(+) cells. STING activation was also highest at two hours with the most pronounced HPV-mediated suppression occurring between one and two hours.

Our data indicates that the slight suppression of STING is both E6 and E7 dependent. In the overexpression system, cells co-expressing both E6 and E7 showed a significant suppression of STING activation compared to vector control cells and cells expressing either oncogene alone. In the TTL cell lines, cells harboring the HPV18 genome exhibited reduced peak pSTING compared to uninfected cells and cells lacking functional E6 and E7. Cells expressing either oncogene alone also showed muted STING activation, though not to the same extent as cells with the full HPV18 genome. These data suggest that while both E6 and E7 are required in an overexpression context, in the context of the full genome, expression of a single oncogene may be sufficient to suppress STING. Recent research has identified that E5 can modulate the cGAS/STING pathway (Miyauchi et al., 2023) and thus may help suppress STING in the TTL cell lines. Further, while the TTL cell lines demonstrated that expression of either oncogene was sufficient for reducing STING phosphorylation, expression of the full genome resulted in the most significant decrease, which highlights a likely combinatorial effect of the HPV oncogenes of STING suppression. This result is similar to what was observed by Lo Cigno et al., who found that knockout of both E6 and E7 had a significant increase on pSTING levels following poly(dA:dT) stimulation (Lo Cigno et al., 2020).

Our data suggests that HPV selectively modulates the transcriptional response downstream of the cGAS/STING pathway. Approximately one third of the genes induced by cGAMP in uninfected cells were suppressed by HPV. These genes were enriched for cytokine activity and antiviral defense and contained the clusters most impacted by IFNAR blockade. This suggests that HPV is suppressing antiviral genes/cytokines downstream of cGAS/STING, particularly type I IFN-driven ISGs. Genes that were induced by cGAMP and were further elevated by HPV (clusters 7, 8, 9, and 11) were primarily increased in the HPV(+) cells at baseline and then followed similar kinetics to that of the uninfected cells. These genes were linked to epithelial proliferation and skin development programs as well as non-canonical NF-κB signaling which has been shown to drive cell cycle progression and proliferation (Chen et al., 2001; Cude et al., 2007; Tang et al., 2008).

The genes in clusters 10 and 12 were elevated at baseline in the HPV(+) cells and then maintained at consistently high levels following cGAMP stimulation while the HPV(-) cells continued to show a dynamic response. This persistent elevation and lack of responsiveness may be indicative of HPV-mediated epigenetic remodeling. Several studies have demonstrated that HPV reprograms chromatin and histone landscapes to modulate host immune responses. For example, Lo Cigno et al. showed that HPV16 E7 induced expression of the histone methyltransferase SUV39H1, which silenced the promotors of cGAS, STING, and RIG-I thereby reducing type I IFN production (Lo Cigno et al., 2020). Similarly, HPV16 and HPV18 have been reported to increase repressive H3K27me3 marks on the promotors of innate immune genes, resulting in a suppression of type I IFNs (Albertini et al., 2018). HPV can also epigenetically enhance gene expression, as demonstrated by Zhang et al., who showed that HPV16 E7 increased histone acetylation at E2F1 promotors to generate transcriptionally active chromatin and promote cell cycle progression (Zhang et al., 2004). Collectively, these observations suggest that HPV may epigenetically elevate baseline expression of select genes, such as those related to epithelial proliferation, while limiting further induction by IRF3/NF-κB.

Beyond epigenetic regulation, other mechanisms may explain the altered signaling output in HPV(+) cells. One possibility is that the increase in cGAMP may serve to exhaust the pathway. A recent study by Li et al. found that chronic STING stimulation diminished STING responsiveness. Repeated cGAMP exposure resulted in an “exhausted” STING phenotype with decrease in STING activation and downstream IFN signaling but increased ER-stress response and NF-κB target genes (Li et al., 2023). The increase in cGAMP we observed could therefore represent a mechanism by which HPV works to “exhaust” STING signaling. Consistent with this idea, we observed a decrease in pSTING and pIRF3 as well as an increase in non-canonical NF-κB signaling.

Another possibility is that the slight suppression of IRF3 results in a change in transcription factor signaling dynamics. Prior studies of signaling dynamics have shown that signal transduction pathways, including cGAS/STING, can exhibit threshold-dependent behavior where minor upstream changes lead to pronounced differences in gene expression (Cheng et al., 2021; Kabelitz et al., 2022; Son et al., 2022; Zinkle & Mohammadi, 2018). For example, Zhou et al. demonstrated that differing levels of pIRF3 drove distinct downstream gene expression patterns (Zhou et al., 2020). Thus, the modest decrease in pIRF3 we observed in HPV(+) cells may shift transcription factor activity away from a type I IFN response and towards more favorable downstream signaling for HPV.

In summary, our data suggests that HPV does not simply turn the cGAS/STING pathway “off,” but rather reshapes its signaling output. This altered output may arise from pathway exhaustion (Li et al., 2023), changes in transcription factor dynamics that modify downstream gene induction thresholds (Zhou et al., 2020), or epigenetic remodeling that renders antiviral promoters less responsive to activation (Albertini et al., 2018; Gao et al., 2021; Lo Cigno et al., 2020; Zhang et al., 2004). Collectively, these findings indicate that E6 and E7 are required and sufficient to increase cGAMP levels and suppress STING activation and that HPV reshapes activation dynamics downstream of the cGAS/STING pathway in a way that blunts antiviral responses and may favor persistence.

## Supplementary Data

Table S1. Genes upregulated by cGAMP and increased by HPV. 284 genes that are significantly upregulated in HPV(+) versus HPV(−) keratinocytes at one or more time points following cGAMP stimulation (3 hr, 6 hr, 9 hr) and that also overlap with genes significantly upregulated by cGAMP in HPV(−) cells at any time point post stimulation (0hr vs 3hr, 6hr, or 9hr). Significance was set as padj <0.05 and log₂ fold-change ≥ 0.5 at one or more time points.

Table S2. Genes upregulated by cGAMP but suppressed by HPV. 263 genes that are significantly downregulated in HPV(+) versus HPV(−) keratinocytes at one or more time points following cGAMP stimulation (3 hr, 6 hr, 9 hr) and that also overlap with genes significantly upregulated by cGAMP in HPV(−) cells at any time point post stimulation (0hr vs 3hr, 6hr, or 9hr). Significance was set as padj <0.05 and log₂ fold-change ≤ −0.5 at one or more time points.

Table S3. Genes upregulated by cGAMP but not significantly changed by HPV. 285 genes that are not significantly differentially regulated in HPV(+) versus HPV(−) keratinocytes at any time point following cGAMP stimulation (3hr, 6hr, or 9hr) and that also overlap with genes that are significantly upregulated by cGAMP in HPV(−) cells at any time point post stimulation (0hr vs 3hr, 6hr, or 9hr). These genes have an adjusted p-value >0.05 and/or an absolute log₂ fold-change greater than −0.5 and less than 0.5 at all time points.

Table S4. Genes list by cluster. 547 genes from Tables S1 and S2 grouped into transcriptional clusters based on hierarchical clustering of variance-stabilized, z-scored, RNA-seq expression data. Each gene is assigned to a cluster (1–12) representing groups of genes with similar expression trajectories.

## Citations

Abe, T., & Barber, G. N. (2014). Cytosolic-DNA-Mediated, STING-Dependent Proinflammatory Gene Induction Necessitates Canonical NF-κB Activation through TBK1. Journal of Virology, 88(10), 5328–5341. 10.1128/JVI.00037-14

Ablasser, A., Schmid-Burgk, J. L., Hemmerling, I., Horvath, G. L., Schmidt, T., Latz, E., & Hornung, V. (2013). Cell intrinsic immunity spreads to bystander cells via the intercellular transfer of cGAMP. Nature, 503(7477), 530–534. 10.1038/nature12640

Albertini, S., Cigno, I. L., Calati, F., Andrea, M. D., Borgogna, C., Dell’Oste, V., Landolfo, S., & Gariglio, M. (2018). HPV18 Persistence Impairs Basal and DNA Ligand–Mediated IFN-β and IFN-λ1 Production through Transcriptional Repression of Multiple Downstream Effectors of Pattern Recognition Receptor Signaling. The Journal of Immunology, 200(6), 2076–2089. 10.4049/jimmunol.1701536

Balka, K. R., Louis, C., Saunders, T. L., Smith, A. M., Calleja, D. J., D’Silva, D. B., Moghaddas, F., Tailler, M., Lawlor, K. E., Zhan, Y., Burns, C. J., Wicks, I. P., Miner, J. J., Kile, B. T., Masters, S. L., & De Nardo, D. (2020). TBK1 and IKKε Act Redundantly to Mediate STING-Induced NF-κB Responses in Myeloid Cells. Cell Reports, 31(1), 107492. 10.1016/j.celrep.2020.03.056

Bartz, S. R., & Vodicka, M. A. (1997). Production of high-titer human immunodeficiency virus type 1 pseudotyped with vesicular stomatitis virus glycoprotein. Methods (San Diego, Calif.), 12(4), 337–342. 10.1006/meth.1997.0487

Bienkowska-Haba, M., Zwolinska, K., Keiffer, T., Scott, R. S., & Sapp, M. (2023). Human Papillomavirus Genome Copy Number Is Maintained by S-Phase Amplification, Genome Loss to the Cytosol during Mitosis, and Degradation in G1 Phase. Journal of Virology, 97(2), e01879–22. 10.1128/jvi.01879-22

Bortnik, V., Wu, M., Julcher, B., Salinas, A., Nikolic, I., Simpson, K. J., McMillan, N. AJ., & Idris, A. (2021). Loss of HPV type 16 E7 restores cGAS-STING responses in human papilloma virus-positive oropharyngeal squamous cell carcinomas cells. Journal of Microbiology, Immunology and Infection, 54(4), 733–739. 10.1016/j.jmii.2020.07.010

Brimer, N., & Vande Pol, S. (2022). Human papillomavirus type 16 E6 induces cell competition. PLoS Pathogens, 18(3), e1010431. 10.1371/journal.ppat.1010431

Carozzi, F., Confortini, M., Dalla Palma, P., Del Mistro, A., Gillio-Tos, A., De Marco, L., Giorgi-Rossi, P., Pontenani, G., Rosso, S., Sani, C., Sintoni, C., Segnan, N., Zorzi, M., Cuzick, J., Rizzolo, R., Ronco, G., & New Technologies for Cervival Cancer Screening (NTCC) Working Group. (2008). Use of p16-INK4A overexpression to increase the specificity of human papillomavirus testing: A nested substudy of the NTCC randomised controlled trial. The Lancet. Oncology, 9(10), 937–945. 10.1016/S1470-2045(08)70208-0

Chapman, S., McDermott, D. H., Shen, K., Jang, M. K., & McBride, A. A. (2014). The effect of Rho kinase inhibition on long-term keratinocyte proliferation is rapid and conditional. Stem Cell Research & Therapy, 5(2), 60. 10.1186/scrt449

Chen, F., Castranova, V., & Shi, X. (2001). New Insights into the Role of Nuclear Factor-κB in Cell Growth Regulation. The American Journal of Pathology, 159(2), 387–397. 10.1016/s0002-9440(10)61708-7

Chen, H., Chen, H., Zhang, J., Wang, Y., Simoneau, A., Yang, H., Levine, A. S., Zou, L., Chen, Z., & Lan, L. (2020). cGAS suppresses genomic instability as a decelerator of replication forks. Science Advances, 6(42), eabb8941. 10.1126/sciadv.abb8941

Chen, Q., Sun, L., & Chen, Z. J. (2016). Regulation and function of the cGAS–STING pathway of cytosolic DNA sensing. Nature Immunology, 17(10), Article 10. 10.1038/ni.3558

Cheng, Q. J., Ohta, S., Sheu, K. M., Spreafico, R., Adelaja, A., Taylor, B., & Hoffmann, A. (2021). NF-κB dynamics determine the stimulus specificity of epigenomic reprogramming in macrophages. Science (New York, N.Y.), 372(6548), 1349–1353. 10.1126/science.abc0269

Chesson, H. W., Dunne, E. F., Hariri, S., & Markowitz, L. E. (2014). The Estimated Lifetime Probability of Acquiring Human Papillomavirus in the United States. Sexually Transmitted Diseases, 41(11), 660–664. 10.1097/OLQ.0000000000000193

Coursey, T. L., & McBride, A. A. (2021). Development of Keratinocyte Cell Lines containing Extrachromosomal Human Papillomavirus Genomes. Current Protocols, 1(9), e235. 10.1002/cpz1.235

Crossley, M. P., Song, C., Bocek, M. J., Choi, J.-H., Kousorous, J., Sathirachinda, A., Lin, C., Brickner, J. R., Bai, G., Lans, H., Vermeulen, W., Abu-Remaileh, M., & Cimprich, K. A. (2023). R-loop-derived cytoplasmic RNA-DNA hybrids activate an immune response. Nature, 613(7942), 187–194. 10.1038/s41586-022-05545-9

Cude, K., Wang, Y., Choi, H.-J., Hsuan, S.-L., Zhang, H., Wang, C.-Y., & Xia, Z. (2007). Regulation of the G2–M cell cycle progression by the ERK5–NFκB signaling pathway. The Journal of Cell Biology, 177(2), 253–264. 10.1083/jcb.200609166

Diner, E. J., Burdette, D. L., Wilson, S. C., Monroe, K. M., Kellenberger, C. A., Hyodo, M., Hayakawa, Y., Hammond, M. C., & Vance, R. E. (2013). The innate immune DNA sensor cGAS produces a noncanonical cyclic dinucleotide that activates human STING. Cell Reports, 3(5), 1355–1361. 10.1016/j.celrep.2013.05.009

Dobin, A., Davis, C. A., Schlesinger, F., Drenkow, J., Zaleski, C., Jha, S., Batut, P., Chaisson, M., & Gingeras, T. R. (2013). STAR: Ultrafast universal RNA-seq aligner. Bioinformatics, 29(1), 15–21. 10.1093/bioinformatics/bts635

D’Souza, G., Tewari, S. R., Troy, T., Webster-Cyriaque, J., Wiley, D. J., Lahiri, C. D., Palella, F. J., Gillison, M. L., Strickler, H. D., Struijk, L., Waterboer, T., Ho, K., Kwait, J., Lazar, J., Weber, K. M., & Fakhry, C. (2024). Oncogenic Oral Human Papillomavirus Clearance Patterns Over 10 years. Cancer Epidemiology, Biomarkers & Prevention : A Publication of the American Association for Cancer Research, Cosponsored by the American Society of Preventive Oncology, 33(4), 516–524. 10.1158/1055-9965.EPI-23-1272

Duensing, S., & Münger, K. (2002). The Human Papillomavirus Type 16 E6 and E7 Oncoproteins Independently Induce Numerical and Structural Chromosome Instability. Cancer Research, 62(23), 7075–7082.

Duensing, S., & Münger, K. (2004). Mechanisms of genomic instability in human cancer: Insights from studies with human papillomavirus oncoproteins. International Journal of Cancer, 109(2), 157–162. 10.1002/ijc.11691

Durbin, J. E., Fernandez-Sesma, A., Lee, C. K., Rao, T. D., Frey, A. B., Moran, T. M., Vukmanovic, S., García-Sastre, A., & Levy, D. E. (2000). Type I IFN modulates innate and specific antiviral immunity. Journal of Immunology (Baltimore, Md.: 1950), 164(8), 4220–4228. 10.4049/jimmunol.164.8.4220

Ewels, P., Magnusson, M., Lundin, S., & Käller, M. (2016). MultiQC: Summarize analysis results for multiple tools and samples in a single report. Bioinformatics, 32(19), 3047– 3048. 10.1093/bioinformatics/btw354

Fitzgerald, K. A., McWhirter, S. M., Faia, K. L., Rowe, D. C., Latz, E., Golenbock, D. T., Coyle, A. J., Liao, S.-M., & Maniatis, T. (2003). IKKε and TBK1 are essential components of the IRF3 signaling pathway. Nature Immunology, 4(5), Article 5. 10.1038/ni921

Gao, P., Ascano, M., Wu, Y., Barchet, W., Gaffney, B. L., Zillinger, T., Serganov, A. A., Liu, Y., Jones, R. A., Hartmann, G., Tuschl, T., & Patel, D. J. (2013). Cyclic [G(2’,5’)pA(3’,5’)p] is the metazoan second messenger produced by DNA-activated cyclic GMP-AMP synthase. Cell, 153(5), 1094–1107. 10.1016/j.cell.2013.04.046

Gao, Z., Li, W., Mao, X., Huang, T., Wang, H., Li, Y., Liu, B., Zhong, J., Renjie, C., Jin, J., & Li, Y. (2021). Single-nucleotide methylation specifically represses type I interferon in antiviral innate immunity. The Journal of Experimental Medicine, 218(3), e20201798. 10.1084/jem.20201798

Gui, X., Yang, H., Li, T., Tan, X., Shi, P., Li, M., Du, F., & Chen, Z. J. (2019). Autophagy Induction via STING Trafficking Is a Primordial Function of the cGAS Pathway. Nature, 567(7747), 262–266. 10.1038/s41586-019-1006-9

Gusho, E., & Laimins, L. A. (2022). Human papillomaviruses sensitize cells to DNA damage induced apoptosis by targeting the innate immune sensor cGAS. PLOS Pathogens, 18(7), e1010725. 10.1371/journal.ppat.1010725

Halbert, C. L., Demers, G. W., & Galloway, D. A. (1991). The E7 gene of human papillomavirus type 16 is sufficient for immortalization of human epithelial cells. Journal of Virology, 65(1), 473–478. 10.1128/jvi.65.1.473-478.1991

Hopfner, K.-P., & Hornung, V. (2020). Molecular mechanisms and cellular functions of cGAS– STING signalling. Nature Reviews Molecular Cell Biology, 21(9), Article 9. 10.1038/s41580-020-0244-x

Ishikawa, H., & Barber, G. N. (2008). STING is an endoplasmic reticulum adaptor that facilitates innate immune signalling. Nature, 455(7213), 674–678. 10.1038/nature07317

Ishikawa, H., Ma, Z., & Barber, G. N. (2009). STING regulates intracellular DNA-mediated, type I interferon-dependent innate immunity. Nature, 461(7265), Article 7265. 10.1038/nature08476

Kabelitz, D., Zarobkiewicz, M., Heib, M., Serrano, R., Kunz, M., Chitadze, G., Adam, D., & Peters, C. (2022). Signal strength of STING activation determines cytokine plasticity and cell death in human monocytes. Scientific Reports, 12, 17827. 10.1038/s41598-022-20519-7

Karin, M. (1999). How NF-κB is activated: The role of the IκB kinase (IKK) complex. Oncogene, 18(49), Article 49. 10.1038/sj.onc.1203219

Korzeniewski, N., Spardy, N., Duensing, A., & Duensing, S. (2011). Genomic Instability and Cancer: Lessons Learned from Human Papillomaviruses. Cancer Letters, 305(2), 113–122. 10.1016/j.canlet.2010.10.013

Lau, L., Gray, E. E., Brunette, R. L., & Stetson, D. B. (2015). DNA tumor virus oncogenes antagonize the cGAS-STING DNA-sensing pathway. Science, 350(6260), 568–571. 10.1126/science.aab3291

Li, J., Hubisz, M. J., Earlie, E. M., Duran, M. A., Hong, C., Varela, A. A., Lettera, E., Deyell, M., Tavora, B., Havel, J. J., Phyu, S. M., Amin, A. D., Budre, K., Kamiya, E., Cavallo, J.-A., Garris, C., Powell, S., Reis-Filho, J. S., Wen, H., … Bakhoum, S. F. (2023). Non-cell-autonomous cancer progression from chromosomal instability. Nature, 620(7976), 1080–1088. 10.1038/s41586-023-06464-z

Liu, H., Zhang, H., Wu, X., Ma, D., Wu, J., Wang, L., Jiang, Y., Fei, Y., Zhu, C., Tan, R., Jungblut, P., Pei, G., Dorhoi, A., Yan, Q., Zhang, F., Zheng, R., Liu, S., Liang, H., Liu, Z., … Ge, B. (2018). Nuclear cGAS suppresses DNA repair and promotes tumorigenesis. Nature, 563(7729), 131–136. 10.1038/s41586-018-0629-6

Liu, T., Zhang, L., Joo, D., & Sun, S.-C. (2017). NF-κB signaling in inflammation. Signal Transduction and Targeted Therapy, 2(1), 1–9. 10.1038/sigtrans.2017.23

Lo Cigno, I., Calati, F., Borgogna, C., Zevini, A., Albertini, S., Martuscelli, L., De Andrea, M., Hiscott, J., Landolfo, S., & Gariglio, M. (2020). Human Papillomavirus E7 Oncoprotein Subverts Host Innate Immunity via SUV39H1-Mediated Epigenetic Silencing of Immune Sensor Genes. Journal of Virology, 94(4), e01812–19. 10.1128/JVI.01812-19

Love, M. I., Huber, W., & Anders, S. (2014). Moderated estimation of fold change and dispersion for RNA-seq data with DESeq2. Genome Biology, 15(12), 550. 10.1186/s13059-014-0550-8

Luo, X., Donnelly, C. R., Gong, W., Heath, B. R., Hao, Y., Donnelly, L. A., Moghbeli, T., Tan, Y. S., Lin, X., Bellile, E., Kansy, B. A., Carey, T. E., Brenner, J. C., Cheng, L., Polverini, P. J., Morgan, M. A., Wen, H., Prince, M. E., Ferris, R. L., … Lei, Y. L. (2020). HPV16 drives cancer immune escape via NLRX1-mediated degradation of STING. The Journal of Clinical Investigation, 130(4), 1635–1652. 10.1172/JCI129497

Luo, Y., Niu, M., Liu, Y., Zhang, M., Deng, Y., Mu, D., Xu, J., & Hong, S. (2024). Oncoproteins E6 and E7 upregulate topoisomerase I to activate the cGAS-PD-L1 pathway in cervical cancer development. Frontiers in Pharmacology, 15, 1450875. 10.3389/fphar.2024.1450875

McLaughlin-Drubin, M. E., Park, D., & Munger, K. (2013). Tumor suppressor p16INK4A is necessary for survival of cervical carcinoma cell lines. Proceedings of the National Academy of Sciences of the United States of America, 110(40), 16175–16180. 10.1073/pnas.1310432110

Mehta, K., & Laimins, L. (2018). Human Papillomaviruses Preferentially Recruit DNA Repair Factors to Viral Genomes for Rapid Repair and Amplification. mBio, 9(1), e00064–18. 10.1128/mBio.00064-18

Meyers, C., Frattini, M. G., Hudson, J. B., & Laimins, L. A. (1992). Biosynthesis of Human Papillomavirus from a Continuous Cell Line Upon Epithelial Differentiation. Science, 257(5072), 971–973. 10.1126/science.1323879

Miyauchi, S., Kim, S. S., Jones, R. N., Zhang, L., Guram, K., Sharma, S., Schoenberger, S. P., Cohen, E. E. W., Califano, J. A., & Sharabi, A. B. (2023). Human papillomavirus E5 suppresses immunity via inhibition of the immunoproteasome and STING pathway. Cell Reports, 42(5), 112508. 10.1016/j.celrep.2023.112508

Mukai, K., Konno, H., Akiba, T., Uemura, T., Waguri, S., Kobayashi, T., Barber, G. N., Arai, H., & Taguchi, T. (2016). Activation of STING requires palmitoylation at the Golgi. Nature Communications, 7(1), Article 1. 10.1038/ncomms11932

Saitoh, T., Fujita, N., Hayashi, T., Takahara, K., Satoh, T., Lee, H., Matsunaga, K., Kageyama, S., Omori, H., Noda, T., Yamamoto, N., Kawai, T., Ishii, K., Takeuchi, O., Yoshimori, T., & Akira, S. (2009). Atg9a controls dsDNA-driven dynamic translocation of STING and the innate immune response. Proceedings of the National Academy of Sciences, 106(49), 20842–20846. 10.1073/pnas.0911267106

Samuel, C. E. (2001). Antiviral Actions of Interferons. Clinical Microbiology Reviews, 14(4), 778–809. 10.1128/CMR.14.4.778-809.2001

Saulters, E. L., Kennedy, P. T., Carter, R. J., Alsufyani, A., Jones, T. M., Woolley, J. F., & Dahal, L. N. (2024). Differential Regulation of the STING Pathway in Human Papillomavirus–Positive and -Negative Head and Neck Cancers. Cancer Research Communications, 4(1), 118–133. 10.1158/2767-9764.CRC-23-0299

Scagnolari, C., Cannella, F., Pierangeli, A., Mellinger Pilgrim, R., Antonelli, G., Rowley, D., Wong, M., Best, S., Xing, D., Roden, R. B. S., & Viscidi, R. (2020). Insights into the Role of Innate Immunity in Cervicovaginal Papillomavirus Infection from Studies Using Gene-Deficient Mice. Journal of Virology, 94(12), 10.1128/jvi.00087-20. https://doi.org/10.1128/jvi.00087-20

Schneider, C. A., Rasband, W. S., & Eliceiri, K. W. (2012). NIH Image to ImageJ: 25 years of image analysis. Nature Methods, 9(7), 671–675. 10.1038/nmeth.2089

Seavey, S. E., Holubar, M., Saucedo, L. J., & Perry, M. E. (1999). The E7 Oncoprotein of Human Papillomavirus Type 16 Stabilizes p53 through a Mechanism Independent of p19ARF. Journal of Virology, 73(9), 7590–7598. 10.1128/jvi.73.9.7590-7598.1999

Shaikh, M. H., Bortnik, V., McMillan, N. AJ., & Idris, A. (2019). cGAS-STING responses are dampened in high-risk HPV type 16 positive head and neck squamous cell carcinoma cells. Microbial Pathogenesis, 132, 162–165. 10.1016/j.micpath.2019.05.004

Shi, H.-J., Song, H., Zhao, Q.-Y., Tao, C.-X., Liu, M., & Zhu, Q.-Q. (2018). Efficacy and safety of combined high-dose interferon and red light therapy for the treatment of human papillomavirus and associated vaginitis and cervicitis: A prospective and randomized clinical study. Medicine, 97(37), e12398. 10.1097/MD.0000000000012398

Shin, J. R., Avilov, I., Schelhaas, M., Gallardo, F., & McBride, A. A. (2025). Tracking replicating HPV genomes in proliferating keratinocytes. mBio, 0(0), e01308–25. 10.1128/mbio.01308-25

Son, M., Frank, T., Holst-Hansen, T., Wang, A. G., Junkin, M., Kashaf, S. S., Trusina, A., & Tay, S. (2022). Spatiotemporal NF-κB dynamics encodes the position, amplitude, and duration of local immune inputs. Science Advances, 8(35), eabn6240. 10.1126/sciadv.abn6240

Sterling, J., Stanley, M., Gatward, G., & Minson, T. (1990). Production of human papillomavirus type 16 virions in a keratinocyte cell line. Journal of Virology, 64(12), 6305–6307. 10.1128/JVI.64.12.6305-6307.1990

Sun, W., Li, Y., Chen, L., Chen, H., You, F., Zhou, X., Zhou, Y., Zhai, Z., Chen, D., & Jiang, Z. (2009). ERIS, an endoplasmic reticulum IFN stimulator, activates innate immune signaling through dimerization. Proceedings of the National Academy of Sciences of the United States of America, 106(21), 8653–8658. 10.1073/pnas.0900850106

Tanaka, Y., & Chen, Z. J. (2012). STING Specifies IRF3 Phosphorylation by TBK1 in the Cytosolic DNA Signaling Pathway. Science Signaling, 5(214), ra20–ra20. 10.1126/scisignal.2002521

Tang, R., Zheng, X.-L., Callis, T. E., Stansfield, W. E., He, J., Baldwin, A. S., Wang, D.-Z., & Selzman, C. H. (2008). Myocardin inhibits cellular proliferation by inhibiting NF-κB(p65)-dependent cell cycle progression. Proceedings of the National Academy of Sciences of the United States of America, 105(9), 3362–3367. 10.1073/pnas.0705842105

Templeton, C. W., & Laimins, L. A. (2023). P53-dependent R-loop formation and HPV pathogenesis. Proceedings of the National Academy of Sciences of the United States of America, 120(35), e2305907120. 10.1073/pnas.2305907120

Templeton, C. W., & Laimins, L. A. (2024). HPV induced R-loop formation represses innate immune gene expression while activating DNA damage repair pathways. PLOS Pathogens, 20(8), e1012454. 10.1371/journal.ppat.1012454

Terenzi, F., Saikia, P., & Sen, G. C. (2008). Interferon-inducible protein, P56, inhibits HPV DNA replication by binding to the viral protein E1. The EMBO Journal, 27(24), 3311–3321. 10.1038/emboj.2008.241

Tolbert, E., Dacus, D., Pollina, R., & Wallace, N. A. (2024). Cutaneous human papillomavirus E6 impairs the cGAS-STING pathway. bioRxiv: The Preprint Server for Biology, 2024.11.29.625575. 10.1101/2024.11.29.625575

Uhlorn, B. L., Jackson, R., Li, S., Bratton, S. M., Van Doorslaer, K., & Campos, S. K. (2020). Vesicular trafficking permits evasion of cGAS/STING surveillance during initial human papillomavirus infection. PLoS Pathogens, 16(11), e1009028. 10.1371/journal.ppat.1009028

Van Doorslaer, K., Porter, S., McKinney, C., Stepp, W. H., & McBride, A. A. (2016). Novel recombinant papillomavirus genomes expressing selectable genes. Scientific Reports, 6(1), Article 1. 10.1038/srep37782

Wieland, S. F., Guidotti, L. G., & Chisari, F. V. (2000). Intrahepatic induction of alpha/beta interferon eliminates viral RNA-containing capsids in hepatitis B virus transgenic mice. Journal of Virology, 74(9), 4165–4173. 10.1128/jvi.74.9.4165-4173.2000

Xu, S., Hu, E., Cai, Y., Xie, Z., Luo, X., Zhan, L., Tang, W., Wang, Q., Liu, B., Wang, R., Xie, W., Wu, T., Xie, L., & Yu, G. (2024). Using clusterProfiler to characterize multiomics data. Nature Protocols, 19(11), 3292–3320. 10.1038/s41596-024-01020-z

Zhang, B., Laribee, R. N., Klemsz, M. J., & Roman, A. (2004). Human papillomavirus type 16 E7 protein increases acetylation of histone H3 in human foreskin keratinocytes. Virology, 329(1), 189–198. 10.1016/j.virol.2004.08.009

Zhao, B., Xu, P., Rowlett, C. M., Jing, T., Shinde, O., Lei, Y., West, A. P., Liu, W. R., & Li, P. (2020). The molecular basis of tight nuclear tethering and inactivation of cGAS. Nature, 587(7835), 673–677. 10.1038/s41586-020-2749-z

Zheng, J., Mo, J., Zhu, T., Zhuo, W., Yi, Y., Hu, S., Yin, J., Zhang, W., Zhou, H., & Liu, Z. (2020). Comprehensive elaboration of the cGAS-STING signaling axis in cancer development and immunotherapy. Molecular Cancer, 19, 133. 10.1186/s12943-020-01250-1

Zhong, B., Yang, Y., Li, S., Wang, Y.-Y., Li, Y., Diao, F., Lei, C., He, X., Zhang, L., Tien, P., & Shu, H.-B. (2008). The adaptor protein MITA links virus-sensing receptors to IRF3 transcription factor activation. Immunity, 29(4), 538–550. 10.1016/j.immuni.2008.09.003

Zhou, J., Sun, T., Jin, S., Guo, Z., & Cui, J. (2020). Dual Feedforward Loops Modulate Type I Interferon Responses and Induce Selective Gene Expression during TLR4 Activation. iScience, 23(2), 100881. 10.1016/j.isci.2020.100881

Zinkle, A., & Mohammadi, M. (2018). A threshold model for receptor tyrosine kinase signaling specificity and cell fate determination. F1000Research, 7, F1000 Faculty Rev-872. 10.12688/f1000research.14143.1

